# From form to function: morphology as a proxy for fish life histories

**DOI:** 10.1101/2025.11.12.688009

**Authors:** Victoria Dixon, Isabel M. Smallegange

**Author notes:** Corresponding author: Isabel Smallegange.

## Abstract

1. Life history strategies emerge from trade-offs between growth, survival and reproduction and are key predictors of how populations respond to environmental disturbances. However, estimating these strategies typically requires detailed demographic data, which are unavailable for many species. Because morphological traits govern whole-organism performance, selection on ecological performance can link morphology with life history strategies. Morphological traits can thus be practical proxies for life history strategies, offering a scalable approach for data-deficient populations.
2. To test the hypothesis that morphological traits predict major life-history axes via shared performance trade-offs, we parameterised dynamic energy budget integral projection models for 290 marine and freshwater fish species to quantify life history strategies and measured eight morphological traits from lateral-view photographs for each species. We used phylogenetically corrected principal component analysis to summarise life history strategies and morphological traits, and tested whether morphology predicts life history strategies.
3. Lateral size morphology, comprising body elongation, relative eye size and oral gape position predicted generation turnover depending on water column position, and predicted reproductive output depending on clade.
4. Our results support the hypothesis that morphologies linked to ecological performance scale up to shape demographic strategies, providing proof of concept that morphology can predict life-history strategies. They also highlight the potential to develop performance-based trait proxies for rapid, low-cost estimations of demographic vulnerability and recovery potential across data-poor fish populations—expanding the scope of life-history frameworks for fisheries management and conservation under increasing pressures from overfishing, habitat loss and climate change.

## 1 INTRODUCTION

Life history strategies shape how species grow, survive, and reproduce—ultimately determining how they persist and adapt in a changing world (Stearns, 1992, 2000). These strategies emerge from fundamental trade-offs in allocation of limited resources among growth, survival and reproduction, shaped by evolutionary history and ecological context (Gaillard et al. 2016). Understanding where species fall along major life history axes provides powerful insights into their vulnerability to disturbance, potential for recovery, and long-term persistence (Salguero-Gómez 2017) — issues that are increasingly critical as global conditions change.

Across many taxa, the dominant axis of life history variation corresponds to the *fast–slow* continuum (Gaillard, 1989; Bielby et al., 2007; Salguero-Gómez et al., 2016; Bakewell et al., 2020, but see Lucas et al. 2025). Along the fast–slow continuum, slow-lived species mature late, reproduce less frequently and live longer, whereas fast-lived species reproduce early and often but have shorter lifespans (Pianka, 1970; Sæther, 1987). Many comparative studies also identify a second major axis of life-history variation, but its biological interpretation is less consistent. Depending on the taxa and traits analysed, this secondary axis has been interpreted as an altricial–precocial continuum (Stearns, 1983), a trade-off between offspring number and offspring quality (Bielby et al., 2007), or more generally as variation in reproductive or developmental strategies. Species’ positions along these continua have been used to predict their sensitivity to environmental change, invasions and extinction risk (Morrison and Hero, 2003; Salguero-Gómez, 2017; Paniw et al., 2018; Ozgul et al., 2024). Fast-lived species, for instance, are often assumed to recover more quickly from disturbance due to faster generation turnover, though this relationship is not universal (Rademaker, van Leeuwen and Smallegange, 2024). Nevertheless, such frameworks have become increasingly relevant for conservation, climate impact assessment and fisheries management (King and McFarlane, 2003; Carruthers et al., 2014; Pennino, Coll and Cerviño, 2023; Smallegange, Edwards and Attle, 2025).

The trade-offs that underpin life history strategies also manifest in morphological, behavioural and physiological traits (Reznick, Bryga and Endler, 1990; Violle et al., 2007). If these traits could reliably reflect a species’ life history strategy, they would offer a scalable and cost-effective pathway to infer ecological performance, since functional trait data are easier to collect than demographic time series. Morphological traits are particularly promising because many remain stable throughout an individual’s adult life, enabling robust cross-species comparisons. In plants, morphology–life-history links have been established at broad taxonomic scales, where traits related to stature, leaf economics and reproductive architecture mediate growth and survival trade-offs (Milla et al., 2008; Adler et al., 2013; Salguero-Gómez, 2017). But similar large-scale analyses are rare for other taxa (Stott et al., 2024). Morphological traits are directly linked to whole-organism performance (Lailvaux & Husak, 2014; Lailvaux & Husak, 2017), including locomotion efficiency, foraging success, predator avoidance and energy expenditure (Gatz 1979; Reznick et al. 1990; Violle et al. 2007). Because performance mediates survival, growth and reproduction, selection on ecological function should create consistent evolutionary associations between form and life-history strategy (Clements & Ozgul 2016; Laughlin et al. 2020). In fish, for example, traits that influence swimming hydrodynamics, visual foraging and feeding mechanics – such as body elongation, eye size and oral gape position – are therefore expected to covary with generation time, reproductive output and life-history pace, particularly across environmental gradients where hydrodynamic and foraging-related selective pressures differ (Chea et al. 2021; Webb, 1984). But despite extensive work linking functional traits to demography in terrestrial taxa, particularly plants (Milla et al. 2008; Adler et al. 2013; Salguero-Gómez et al. 2017), large-scale tests of these links in fishes, or other aquatic taxa, remain rare.

This study tests the hypothesis that fish morphological traits, linked to swimming behaviour, food acquisition and predator avoidance, predict major life-history axes via shared performance trade-offs. Their extraordinary morphological diversity arises from both ecological specialisation and strong phylogenetic constraints (Gatz, 1979; Salguero-Gómez et al., 2018), and morphological measurements can now be efficiently extracted from widely available lateral-view photographs (Brosse and Charpin, 2021). Yet, no study has systematically tested whether fish morphology predicts life history strategies across large phylogenetic and ecological scales. Moreover, fish life histories do not always conform to terrestrial models: slow-lived fish species often produce more, not fewer, offspring than fast-lived species (Jeschke and Kokko, 2009; Capdevila et al., 2020), and, when accounting for body size, aquatic species overall exhibit a faster pace of life than terrestrial ones (Capdevila et al., 2020).

To test our hypothesis, we parameterised Dynamic Energy Budget Integral Projection Models (DEB-IPMs; Smallegange et al., 2017) for 290 marine and freshwater species to model survival, growth and reproduction based on species-specific energy budgets and individual-level trade-offs in energy allocation (Kooijman, 2000; Smallegange and Lucas, 2025). From these models, we calculated a set of representative life history traits describing schedules of survival, growth and reproduction, and summarised them using a phylogenetically corrected principal component analysis (PCA) to identify the major axes of fish life history strategies. We applied the same approach to morphological data, using morphological ratios commonly applied in fish biology (Villéger et al., 2017; Brosse and Charpin, 2021) to derive composite morphological axes related to swimming behaviour, food acquisition and predator avoidance. We then assessed the influence of these composite morphological axes, along with phylogenetic ancestry, climate zone and water column position, on species’ positions along the major life history strategy axes, to explore if selection acting on performance differs across environments and lineages. This approach demonstrates a novel application of modelling and analytical techniques to explore the relationship between morphology and life history — providing a pathway for scalable estimation of fish life histories.

## 2 METHODS

### 2.1 Dynamic Energy Budget Integral Projection Model (DEB-IPM)

A DEB-IPM is a demographic model that uses eight life history parameters (Table 1) to capture the demographic rates of survival, growth, reproduction, and the relationship between parent and offspring sizes for females within a population (Smallegange *et al*., 2017; Smallegange and Lucas, 2024). These demographic rates are derived from the simple version of the Kooijman-Metz model (1984). The Kooijman-Metz model assumes that organisms are isomorphic (individuals retain the same overall shape and proportions as they grow, resulting in external morphological similarity, even between individuals with differing phylogenies or biological roles), are born at length *L_b_,* mature at length *L_p_* and ingest food at a rate that is proportional to their experienced energy acquisition level. A feeding level *E*(*Y*) represents the fullness of the animal’s stomach from completely empty (0 ≥ *E(Y)* > 0.1) to completely full (0.9 ≥ *E(Y)* ≥ 1) (Piet and Guruge, 1997). At *E(Y)* = 1, the highest feeding level, individuals grow to their maximum length (*L_m_*) and reproduce at their maximum rate (*R_m_*). As individuals feed and uptake energy, a constant fraction of assimilated energy is allocated to respiration (κ) to cover costs of maintenance and somatic growth. Any remaining energy is allocated to reproduction in adults or development of reproductive organs in juveniles. Note that *R_m_* is proportional to (1 – κ), whereas *L_m_* is proportional to κ, which controls energy conservation. However, the role of κ in a DEB-IPM is mostly implicit, as κ is used as input parameter only in the starvation condition, whereas *R_m_* and *L_m_* are measured directly from data. Growth follows a von Bertalanffy curve characterised by its growth coefficient (ṙ*_B_*). Individuals grow from year *t* to year *t* + 1 if they survive. Their survival chances are determined by mortality rates which differ for adults (*μ_a_*) and juveniles (*μ_j_*). To account for survival during the egg and larval phases, where the highest mortality occurs for ray-finned fish, *R_m_* was multiplied by 0.001 for all actinopterygians (Rademaker, van Leeuwen and Smallegange, 2024). If a species was missing one trait value, either length at birth or reproductive rate, and there was another species of the same genus with a value present, then that value was used. Otherwise, species with missing values were removed from the dataset.

**Table 1:**
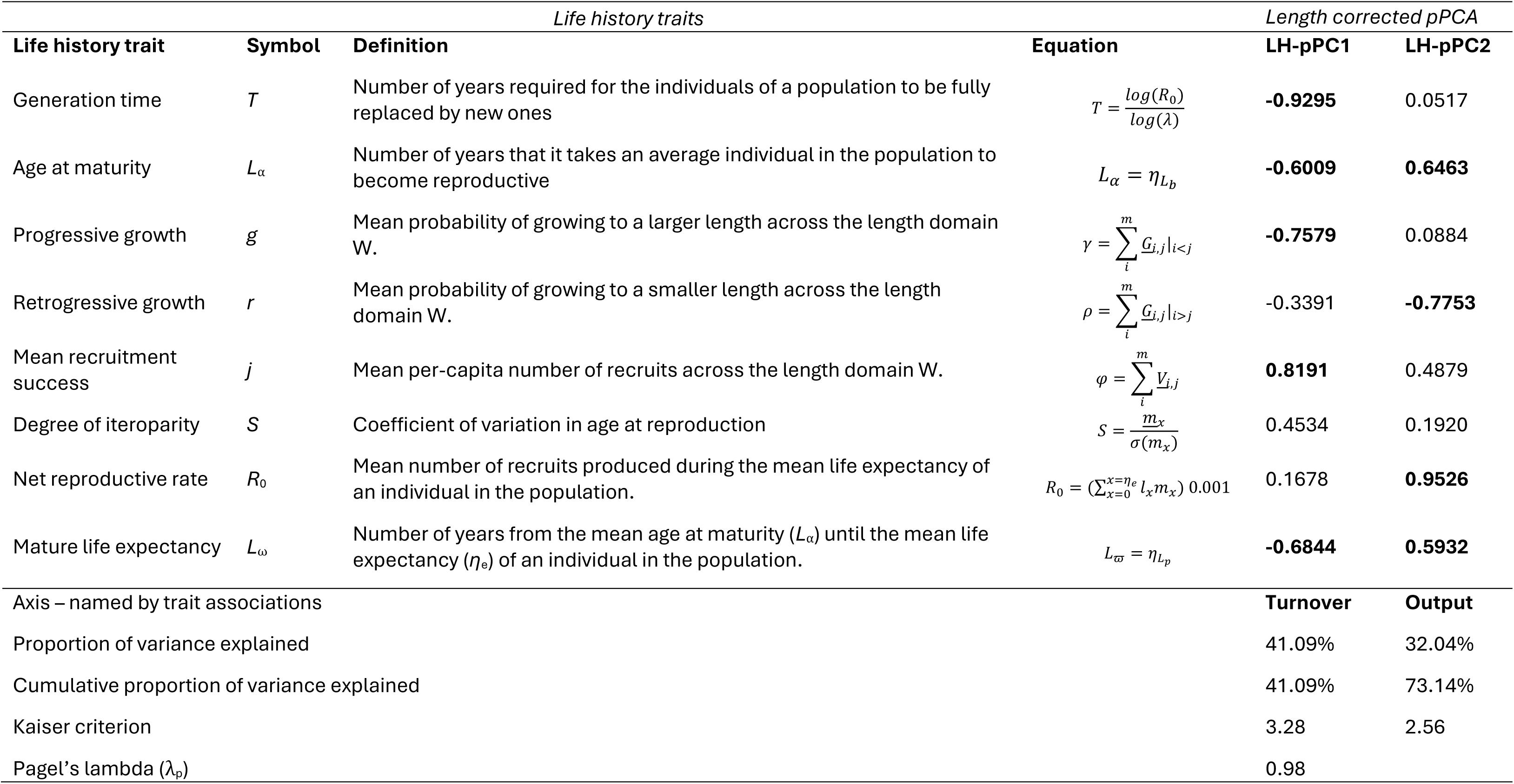
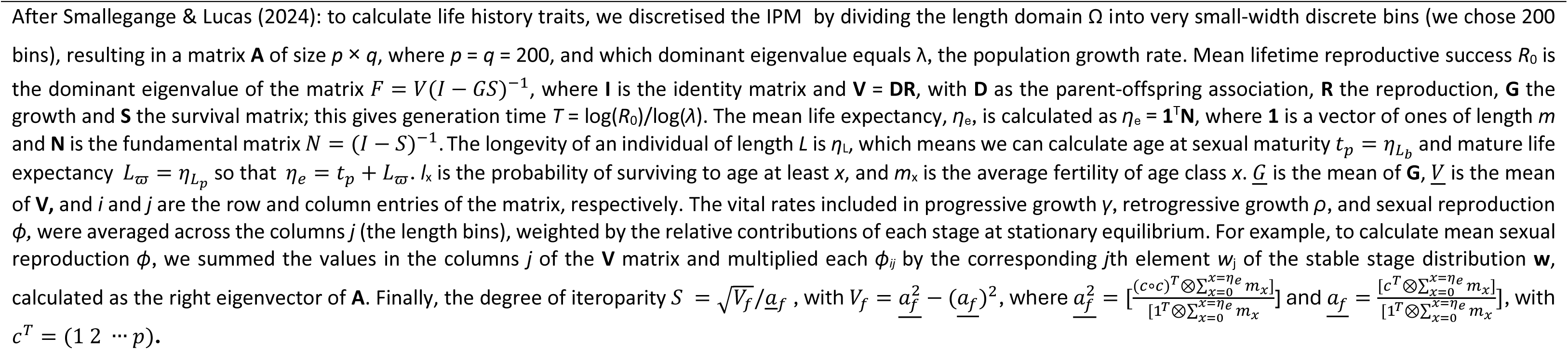
Life history traits calculated from the DEB-IPMs with their definition and how they were calculated. The final two columns are the scores for each of the life history traits on a phylogenetically and length corrected principal component analysis (LH-pPCA). Significant traits for each of the axes (score > 0.5) are in bold. The traits for LH-pPC1 are associated with generation turnover and the traits for LH-pPC2 are associated with reproductive output.

### 2.2 Parameterising the DEB-IPMs

We used the DEBBIES dataset (Smallegange and Lucas, 2024), which contains demographic trait data for 161 ectotherms including 3 actinopterygians and 158 chondrichthyans. We expanded this dataset by retrieving demographic data from FishBase (Froese and Pauly, 2013) and the wider literature (Figure 1). To this end, we developed a filtering method to search through all the entries in FishBase across multiple topics to create a list of species that were likely to have all the necessary demographic data (Appendix A: Filtering Method) (Froese and Pauly, 2013).

**Figure 1.**
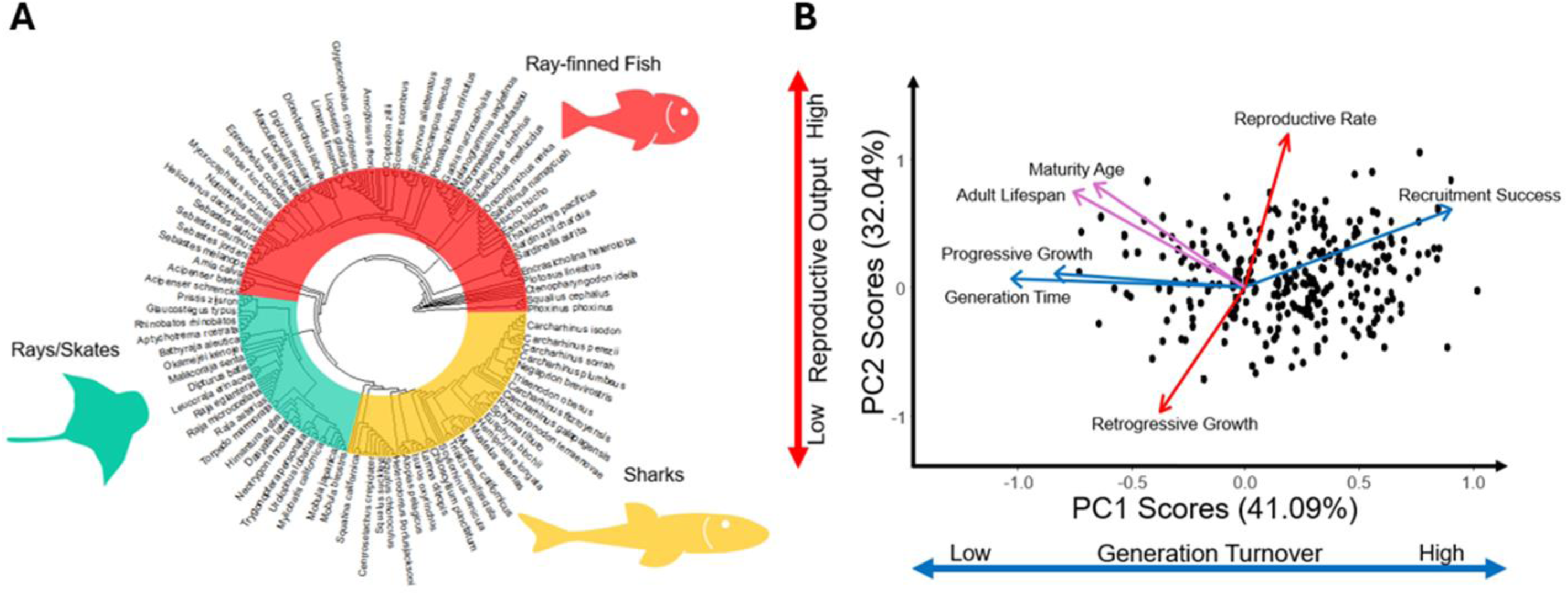
Fish phylogenetic tree and life history biplot. **A**: Circular phylogenetic tree of the 290 study species across three clades: ray-finned fish (red), rays/skates (green), and sharks (yellow). Branch lengths are derived from The Open Tree of Life. A limited number of tip labels are matched to the tree tips given limited space. **B**: A phylogenetically and length corrected PCA biplot of the life histories of 290 fishes (each species is a solid symbol). The strength of the trait loadings onto each of the axes are shown by the length and direction of the black arrows.

#### 2.2.1 Fraction of energy allocated to respiration

The fraction of energy allocated to respiration (κ) values were found on the Add-my-Pet website or, if unavailable, assigned 0.8, which is accepted as a generic value for animals (‘Add-my-pet’, 2023).

#### 2.2.2 Length at birth, at puberty and maximum length

Length at birth for the actinopterygians was initially tested as average egg size, however these values proved too small for the model to run. Instead, the nearest value to egg size for each species was used. Length at puberty (*L_p_*) and maximum length (*L_m_*) were collected from species’ summary pages (Froese and Pauly, 2013). Female length values were preferred over male ones. Typically, measurements were in total length (snout to end of caudal fin) (Table 2). Any values that were in fork length (snout to start of tail fork) or standard length (snout to start of precaudal fin) were converted to total length using the species-specific Length-Length tables from FishBase (Froese and Pauly, 2013). Studies that used a greater sample size were preferred in instances of multiple conversion coefficients.

**Table 2:** Description and diagram of trait measurements for a ray-finned fish taken from lateral view images (adapted from Brosse and Charpin (2021)). Measurements introduced in this study, Caudal fin inner fork distance (CFi) and Caudal fin outer fork distance (CFo) are defined below and highlighted in blue in the figure, along with the other measurements.

| Code | Name | Protocol for measurement |
| --- | --- | --- |
| BL | Body length | Standard length (snout to caudal fin base) |
| Bd | Body depth | Maximum body depth |
| Hd | Head depth | Head depth at the vertical of the eye |
| CPd | Caudal peduncle depth | Minimum depth of the caudal peduncle |
| CFd | Caudal fin depth | Maximum depth of the caudal fin |
| Ed | Eye diameter | Vertical diameter of the eye |
| Eh | Eye position | Vertical distance between the centre of the eye and the bottom of the body |
| Mo | Mouth height | Vertical distance from the top of the mouth to the bottom of the body |
| JI | Maxillary jaw length | Length from snout to the corner of mouth |
| PFI | Pectoral fin length | Length of the longest ray of the pectoral fin |
| PFj | Pectoral fin position | Vertical distance between the upper insertion of the pectoral fin to the bottom of the body |
| CFi | Caudal fin inner fork distance | Beginning of caudal fin (from the lateral line of the spine) to the divide between the upper and lower lobes |
| CFo | Caudal fin outer fork distance | Beginning of caudal fin (from the top of the body) to the furthest tip of the upper lobe |

#### 2.2.3 Adult and juvenile mortality rates

Maximum reported age (*t_max_*) was also collected from the FishBase summary pages for use in mortality calculations (Froese and Pauly, 2013). Mortality rates were calculated using the same method as the DEBBIES dataset for consistency. Adult mortality rate was calculated as 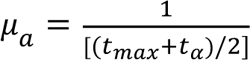 (Pardo *et al*., 2016; Smallegange and Lucas, 2024), where age at maturity (*t_α_*) was calculated as an average of the recorded observations (*t_max_*) for females and unsexed entries from maturity studies pages on FishBase (Froese and Pauly, 2013). For juveniles, 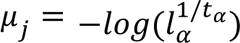, where survival to maturity *l_α_* = *e*^−*μ*^*^a^*^(*t*^*^max^*^−*t*^*^α^*^)^. Unsexed data were included as female-only data were not available for all species. Male or mixed sex data were only used in the absence of other options. For most actinopterygians, the highest proportion of mortalities takes place during the egg and larval stages, so an additional survival factor was introduced into the reproductive rates as described below (Rademaker, van Leeuwen and Smallegange, 2024).

#### 2.2.4 von Bertalanffy growth rate

The von Bertalanffy growth rate coefficient (ṙ*_B_*) was sourced from the species-specific growth parameters available from FishBase (Froese and Pauly, 2013). Some of the FishBase coefficients were derived from equations where the infinite length was more/less than 1/3 of the maximum length – making them less reliable, so using maximum observed age to calculate growth rate, (ṙ*_B_* = −*log*(1 − 0.95)*t_max_*), was tested as an alternative. A one-way analysis of variance (ANOVA) determined that using maximum age to find the coefficients produced significantly different values compared to the reliable growth rate values from FishBase (F_1,222_=4.311, p=0.039), so all the FishBase values were retained for consistency (Froese and Pauly, 2013).

#### 2.2.5 Maximum reproductive rate

Maximum reproductive rate (*R_m_*) was calculated using fecundity data (maximum number of eggs possibly produced across a female’s lifetime) from the species’ reproduction pages on FishBase (Froese and Pauly, 2013). Where multiple observations were recorded, an average of only the maximum values (including single values if a range was not present) was calculated. Maximum fecundity was then divided by the species’ adult lifespan to represent maximum clutch size per year – unless the adult lifespan was less than 1 year then maximum fecundity was left unchanged. To capture egg and larval mortalities, reproductive rate was multiplied by 0.001 for all actinopterygians (Houde and Zastrow, 1993; Rademaker, van Leeuwen and Smallegange, 2024; Smallegange and Lucas, 2024).

#### 2.2.6 Final dataset

The above methods added 160 actinopterygians (21 from Rademaker, van Leeuwen and Smallegange (2024) which included four demographic parameters (*L_p_, L_m_*, ṙ*_B_* and *R_m_*), the others were collected following the protocols above) to the DEBBIES dataset totalling 321 species (158 chondrichthyans and 163 actinopterygians; species list in the Supplementary Material).

### 2.3 Quantifying life history strategies

We chose to set the feeding level high at 0.9 across all species for consistency and to better demonstrate where species allocate their energy under realistic optimum conditions (e.g, a feeding level of 0.8 is associated with natural, stable populations in reef manta rays (*Mobula alfredi*) (Smallegange *et al*., 2017).

Using the parameterised DEB-IPMs, we derived eight representative life history traits based on schedules of survival, growth and reproduction: generation time (*T*), age at maturity(*L*_α_), progressive growth (*g*), retrogressive growth (*r*), mean recruitment success (*j*), degree of iteroparity (*S*), net reproductive rate (*R*_0_), and mature life expectancy (*L*_ω_). Details of these variables and how they were calculated can be found in Table 1 (Salguero-Gómez *et al*., 2016; Salguero-Gómez, 2017; Capdevila *et al*., 2020). Analyses were conducted in MATLAB Version 24.2 (The MathWorks Inc, 2024) using the code in Smallegange and Lucas (2024). We also calculated the population growth rate (λ_0_) to test model performance later. To this end, we discretised each DEB-IPM (Smallegange and Lucas, 2024) by dividing the length domain Ω into 200 very small-width discrete bins, resulting in a matrix **A** of size 200×200, of which the dominant eigenvalue equals λ_0_. Finally, we calculated the damping ratio (ξ) with ξ= λ_0_/ λ_2_, where λ_2_ is the highest subdominant eigenvalue of matrix **A** (Caswell, 2001; Capdevila *et al*., 2020). Alternative metrics to capture demographic resilience exist (simulated time-to-recovery or transient growth rates (Stott, Townley and Hodgson, 2011; Capdevila *et al*., 2021), but for ease of comparison with previous studies, we used the damping ratio.

The results of the models were scanned for outliers before proceeding to the next stage of analysis. *Alosa sadipissima* was the only species to produce an NA value and was removed from the dataset. Despite introducing an egg/larval survival factor to reduce reproductive output, several species had abnormally high population growth rates and a high damping ratio. A further 23 species were removed as they met one or both of the following rejection criteria: 0.7 > λ_0_ > 3.0 or ξ > 3.0. The Greenland shark, *Somniosus microcephalus*, was also removed after visual inspection of the data due to its unusually high longevity – a precedent set in other studies (Lucas *et al*., 2025; Smallegange and Lucas, 2024).

The life history traits were log-transformed and scaled (*μ* = 0 and SD = 1) to meet principal component analysis assumptions before use in a phylogenetically corrected and length corrected principal components analysis (pPCA) (Revell, 2012) to identify life history strategy axes. Body length was corrected for in R by calculating the linear model residuals of the log-10 transformed life history traits and maximum body length, previously collected from FishBase, for each species (Gaillard, 1989; Froese and Pauly, 2013; R Core Team, 2024).

Phylogenetic corrections were made at the species level using the *phy.pca* function from the *phytools* library in R (Revell, 2012; Salguero-Gómez *et al*., 2016; Paniw *et al*., 2018; R Core Team, 2024). The phylogenetic tree was generated using the *rotl* package (version 3.1.0) to find each species’ position on the Open Tree of Life (OTT) (Michonneau, Brown and Winter, 2016). The branch lengths of the tree inform on relatedness by scaling proportionally to the divergence times for clades and species. The pPCA linked phylogeny from the tree to the life history traits through a modified covariance matrix. The pPCA also provided an estimate of Pagel’s lambda (λ_p_) – a scaling parameter for the phylogenetic correction between species from phylogenetic independence (0) to the species covarying proportionally to their shared evolutionary history (1) (Pagel, 1999; Freckleton, Harvey and Pagel, 2002). A further four species were excluded because they were not available from the OTT database: *Urobatis halleri*, *Dipturus intermedius, Osmerus dentex,* and *Osmerus mordax*.

PCAs were run with and without the phylogenetic and body length corrections to determine their influence on the structuring of life history strategies. An arbitrary cutoff value of 0.25 was set for λ_p_ as, below it, the impact of their shared evolutionary history was deemed limited (Lucas et al. 2025). The significance of the principal components axes was assessed using Kaiser’s criterion whereby only axes with eigenvalues greater than 1 were retained for further analyses (Kaiser, 1960). This criterion identifies axes that explain more variance than a single original variable and is widely used for dimensionality reduction in multivariate ecological datasets (e.g., Salguero-Gomez et al. 2016; Rademaker et al. 2024; Lucas et al. 2025). Axes with eigenvalues lower than 1 were therefore considered to represent relatively small proportions of variance and were not retained for subsequent analyses.

### 2.4 Collecting morphological trait data

#### 2.4.1 Collection method

Morphological data were collected from lateral view images by following the methodology from FISHMORPH (Brosse and Charpin, 2021). FISHMORPH is a database containing eight unitless ratios (Tables 2, 3) and body length, for 8,432 freshwater fish species (Brosse and Charpin 2021). These ratios are related to swimming efficiency (body elongation, body lateral shape, caudal fin throttling, pectoral fin position and pectoral fin size), prey/predator detection (relative eye size, vertical eye position), and prey capture mechanics (vertical eye position, oral gape position, jaw maxillary length) (Brosse and Charpin, 2021), and contribute to whole-organism performance which mediates survival and reproductive success. Therefore, these performance components are predicted to covary with species’ positions along life-history axes. We calculated these ratios for each of our remaining 290 marine and freshwater fish species from a series of measurements taken from lateral view images (see Appendix B). Rules for unusual morphologies were followed where applicable (Villéger *et al*., 2010; Brosse and Charpin, 2021). FISHMORPH and its protocols were created for actinopterygians, so some adaptations were made to accommodate the chondrichthyans and seahorses into the same methodology (see Appendix B).

**Table 3:**
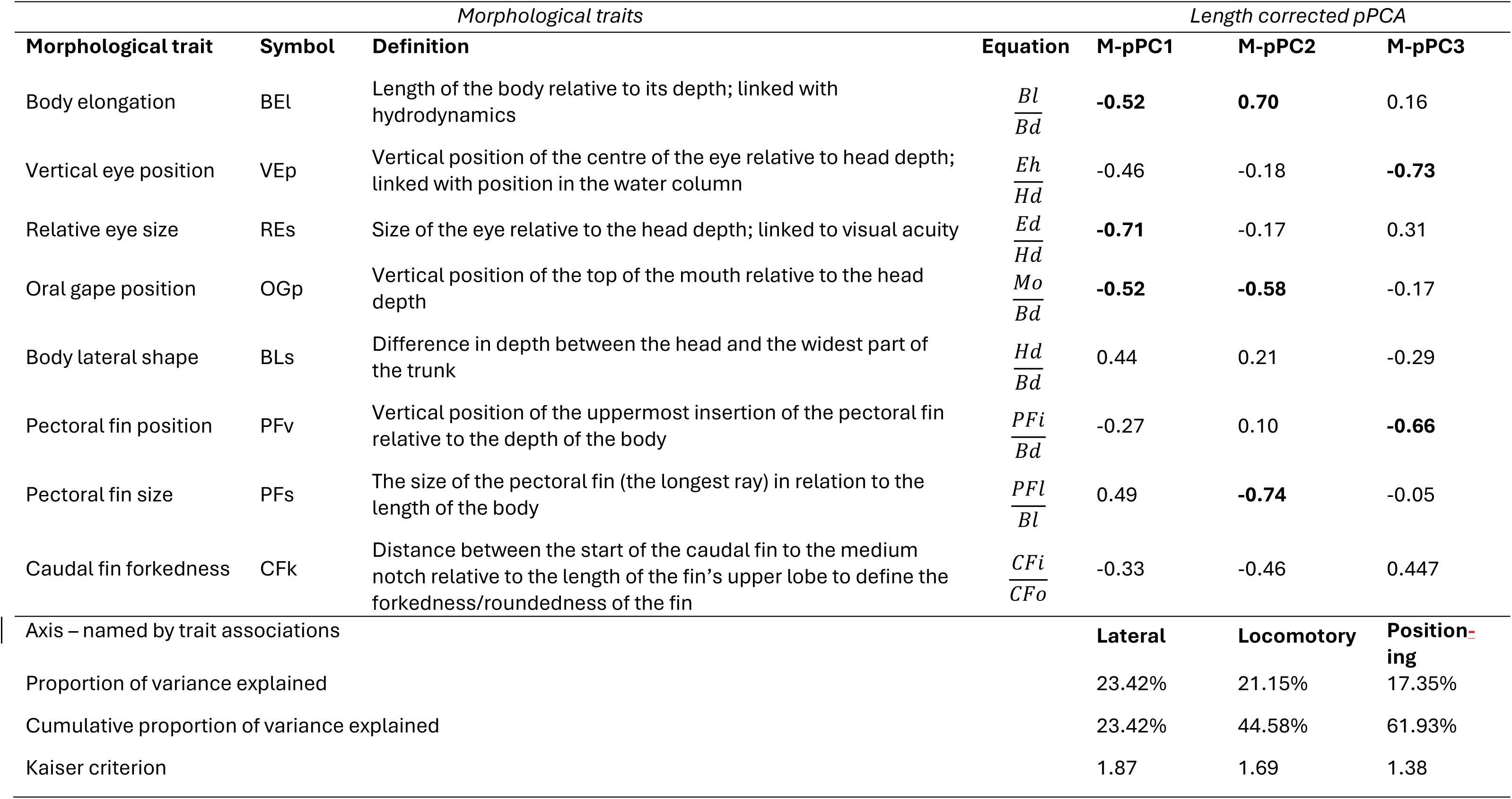
List of the morphological trait ratios included in the morphological PCA, their definitions and how they were calculated (see Table 2). The final three columns show the results of the phylogenetically and length corrected PCA (M-pPCA). Traits that had significant loadings (>0.5) are highlighted in bold. M-pPC axes were named based on the associations of their significant traits: M-pPC1 is related to body size, M-pPC2 is associated with manoeuvring, M-pPC3 is associated with position in the water column relative to prey/predators.

Research grade photographs were found online primarily from iNaturalist (iNaturalist, 2023), FishBase (Froese and Pauly, 2013), institutional websites, or from the personal websites of specialists (image references available in Supplementary Material). All images were available under fair dealing copyright law or had permission given for use in this work. As the life history data is modelled on females of the species, images of females were preferred in cases where extreme sexual dimorphism was known. The largetooth sawfish, *Pristis pristis*, is an exception where a diagram was used instead. Two species had no suitable photographs or diagrams available from any of these sources – *Pristis clavata* and *Sphyrna zaegna*. Their life history data is still available in the raw data file (see Supplementary Material), but they were not included in further analyses.

Images were uploaded to ImageJ 1.54p in a browser and measured in pixels (Schneider, Rasband and Eliceiri, 2012). Ratios were calculated following FISHMORPH (Tables 2, 3) (Brosse and Charpin 2021). Replicates were measured for several species from different pictures to assess intra-specific variability. The resultant ratios for each replicate were averaged and used as the entries for those species.

#### 2.4.2 Introduction of a new morphological ratio

To explore tail shape in more detail, an additional ratio was designed called caudal forkedness (CFk) (Table 3). The fork in caudal fins is related to the individual’s acceleration and manoeuvrability, mediating survival (Mouchet *et al*., 2019). This ratio is calculated by dividing the distance from the start of the caudal fin to the end point that divides the upper and lower lobes (along the lateral line of the spine) by the distance from the start of the caudal fin (along the top of the body) to the tip of the upper lobe (Table 2). Fins that are rounded have a value less than one and fins that have a strong fork have a value greater than one. As this ratio is not in FISHMORPH, and every ratio is required to be related to each other, every species had their values measured from images and not taken directly from the database. In the absence of a caudal fin, CFk was set to zero to represent a pointed tail. Due to the introduction of a new ratio partway through collection, images were entirely re-measured to create data files that included the new caudal fork data. The difference between the former and newer measurements of body length never exceeded fifteen pixels which demonstrates consistency in measurement technique.

### 2.5 Quantifying composite morphological axes

Due to the similarity between some of the traits represented by the morphological ratios, a Pearson’s correlation test was run to determine if ratios covaried before conducting the phylogenetically corrected PCA (see Appendix C). The ratios that had the highest associations with each other were relative maxillary length (RMl) and oral gape position (OGp) as well as caudal forkedness (CFk) and caudal peduncle throttling (CPt). OGp was retained as it was available for more species than RMl. CFk was retained instead of CPt to test the applicability of a new caudal ratio (Mouchet *et al*., 2019). The remaining traits were log-transformed and scaled to meet the PCA assumptions.

We then applied dimension reduction via phylogenetically corrected PCA to extract composite morphological proxies for swimming, foraging and positioning performance, which were then tested against life-history strategy axes. As Pagel’s lambda was high for the life history pPCA, we decided to include phylogenetic corrections in the morphological PCAs as well to identify the main composite morphological axes. The morphological trait data were scaled (*μ* = 0 and SD = 1). The same tree from the life history pPCA was used. Linear model residuals of square root-transformed traits against log-transformed body length were used for the length corrected pPCA, as log-10 transformations were unsuitable for the trait corrections. The axes were assessed using a Kaiser’s criterion (Kaiser, 1960).

The results of the morphological PCA are reported here, as the axes - named according to their strongest trait associations – serve as inputs for the main analyses (see below). Pagel’s lambda was high (λ_p_=0. 99), demonstrating a high phylogenetic signal (Pagel, 1999). A qualitative check demonstrated that trait loadings were different between length corrected pPCA and non-length corrected pPCA so the former was used. The final morphological pPCA was phylogenetically and length corrected. The first three axes were retained from the morphological pPCA because their eigenvalues were higher than unity (1.87, 1.69, and 1.38 respectively), whereas the fourth was not (0.955). The three axes cumulatively explain 61.93% of the total variance (Table 3: M-pPC1 23.42%, M-pPC2 21.15%, and M-pPC3 17.35%). Body elongation (BEl), relative eye size (REs), and oral gape position (OGp) all loaded negatively onto M-pPC1 (Table 3). As M-pPC1 increases, species have deeper bodies, smaller eyes and a lower mouth position relative to their head depth. M-pPC1 is therefore referred to as “*lateral size morphology*”. Body elongation (BEl) loaded negatively onto M-pPC2, but oral gape position (OGp) and pectoral fin size (PFs) loaded positively (Table 3). As M-pPC2 moves from negative to positive scores, species have deeper bodies with larger pectoral fins and a higher mouth position. This trait combination can be associated with acceleration and agility so M-pPC2 will be referred to as “*locomotory morphology*”. For M-pPC3, vertical eye position (VEp) and pectoral fin vertical position (PFv) both loaded negatively (Table 3). As M-pPC3 increases species have eyes in lower positions relative to their heads and lower pectoral fins relative to their trunks. M-pPC3 traits can be associated with position in the water column, relative to their food and predators, and will be referred to as “*positioning morphology*”. Body lateral size and caudal fin forkedness did not load onto the first three axes (Table 3).

### 2.6 Testing which morphological traits predict life history strategy

To test our hypothesis that morphological traits predict major life-history axes, we assessed whether composite morphological axes representing hydrodynamic form (lateral size morphology), locomotor capability (locomotory morphology), and habitat positioning relative to prey and predators (positioning morphology) covaried with species’ positions along the life history axes. This included the generation turnover axis, composed of traits relating to life history speed but none describing reproduction, hence differing from the fast-slow continuum, and a secondary reproductive output axis. We further tested whether these relationships were mediated by ecological context (water column position and climate zone) or phylogenetic ancestry (clade), consistent with predictions that selection acting on performance differs across environments and lineages.

Using data from FishBase, species were divided into broad categories describing the climate zone they are most frequently found in: deep-water (>200m), polar, temperate, and tropical (combined with subtropical) (Froese and Pauly, 2013). Similarly, FishBase was used for species’ most frequented water position: deep-water (>200m), seafloor, pelagic, and reef (Froese and Pauly, 2013). Species’ clades were broadly divided into ray-finned fish, sharks and rays/skates. The influences of climate zone, water position, clade, morphology and their interactions on life history strategies were tested using two General Linear Models (GLMs) using base R (R Core Team, 2024). The response variables were the life history principal component scores of either axis, and the predictor variables were climate zone, water position, clade and each of the morphological principal component (PC) scores (lateral size morphology, locomotory morphology, and positioning morphology). We excluded the interaction between climate zone and water column position because of overlapping categories among deep-water species and insufficient data for certain combinations of these variables (for example, the dataset contained no polar reef species). For both GLMs, the model assumptions of Gaussian errors and homoscedasticity were confirmed by visually inspecting the residual and Q-Q plots in R (R Core Team, 2024). In stepwise regression, non-statistically significant interactions were removed from the models. Higher order interactions were removed first until only statistically significant interactions and fixed factors remained. Significance of interactions were determined by comparing each reduced model with the fuller one it is nested in, using ANOVA tests. Terms were removed if the p-value exceeded 0.05.

## 3 RESULTS

### 3.1 Major life history axes to describe variation in life history traits

After removing outliers or species with missing data, life history traits of 290 species were analysed (Figure 1A). Pagel’s lambda was higher than the cut-off value of 0.25 (λ_p_=0.98), indicating that the phylogenetic signal was strong (Pagel, 1999). A qualitative check of the trait loadings between the pPCA and the length corrected pPCA revealed differences of which axes the life history traits loaded onto. We thus proceeded with the phylogenetically and length corrected pPCA, referred to as the LH-pPCA. Two LH-pPC axes cumulatively explained 73.14% of the total variance in life history traits (Table 1: LH-pPC1 41.09%, LH-pPC2 32.04%), as their eigenvalues were higher than unity (3.287 and 2.563, respectively), whereas the third axis was not (λ = 0.946). Generation time (*T*), mean age at maturity (*L*_α_; note that *L*_α_ is equivalent but not computationally the same as *t*_α_, the observed age at maturity used above to calculate juvenile and adult mortality rates), progressive growth (γ), and mature life expectancy (*L*_ω_) all loaded negatively onto LH-pPC1 whereas mean recruitment success (φ) loaded positively (eigenvalue<0.05) (Table 1). These traits are strongly associated with *generation turnover*. As LH-pPC1 scores increase, our species show shorter lifespans and greater generation turnover (Figure 1B). For LH-pPC2, mean age at maturity (*L*_α_), mature life expectancy (*L*_ω_), and net reproductive rate (*R_0_*) loaded positively whereas retrogressive growth (ρ) loaded negatively onto LH-pPC2 (Table 1). These traits are strongly associated with *reproductive output*. As LH-pPC2 scores move from negative to positive, species produce more offspring (Figure 1B). Degree of iteroparity (*S*) did not load onto either of the first two axes.

### 3.2 Does morphology predict generation turnover?

Consistent with our hypothesis that traits related to swimming and foraging performance covary with life-history speed, lateral size morphology was significantly associated with generation turnover LH-pPC scores, but differently at different water positions (two-way interaction: F_3, 269_=3.3864, p=0.0186). With increasing lateral size morphology, deep-water species had a lower generation turnover whereas species from all other depths had a higher generation turnover (Figure 2). The interaction between water position and clade was also significantly associated with generation turnover (F_6, 269_=2.2039, p=0.0429) Deep-water species had a lower generation turnover than other water positions across all clades, although the difference was most pronounced for the rays/skates (Figure 3A). Climate zone was significantly associated with generation turnover as well (F_3, 269_=9.4796, p<0.001). Deep-water species had statistically significantly lower generation turnover than species in all other climate zones (Figure 3B). Polar species tended to have faster generation turnover than deep-water species but slower than temperate or tropical species (Figure 3B). No other interaction terms were significantly associated with species’ generation turnover: climate zone × positioning morphology (F_2, 243_=0.8474, p=0.4298), climate zone × locomotory morphology (F_3, 245_=1.3128, p=0.2708), climate zone × lateral size morphology (F_3, 248_=0.8677, p=0.4584), water position × positioning morphology (F_3, 251_=0.3993, p=0.7536), water position × locomotory morphology (F_2, 254_=1.3458, p=0.26), clade × positioning morphology (F_2, 257_=0.7117, p=0.4918), clade × locomotory morphology (F_3, 259_=0.3059, p=0.7367), clade × lateral size morphology (F_3, 261_=2.3399, p=0.0983), and clade × climate zone (F_6, 263_=1.937, p=0.0752). Generation turnover was also not significantly associated with locomotory morphology (F_1, 269_=1.6508, p=0.2) or positioning morphology (F_1, 269_=0.6279, p=0.4288). For the other fixed factors, the model became overparametrised, so instead test statistics are reported for lateral size morphology (estimate=1.5913, std. error=0.6816, t=2.335, p=0.0203), water positions including pelagic (estimate=-0.0221, std. error=0.1701, t=-0.130, p=0.8964), reef (estimate=-0.1167, std. error=0.1734, t=-0.673, p=0.5013), and seafloor (estimate=-0.2226, std. error=0.1594, t=-1.397, p=0.1636), as well as the clades: rays/skates (estimate=-0.6308, std. error=0.2985, t=-2.113, p=0.0355) and sharks (estimate=0.0110, std. error=0.1518, t=0.073, p=0.9419).

**Figure 2:**
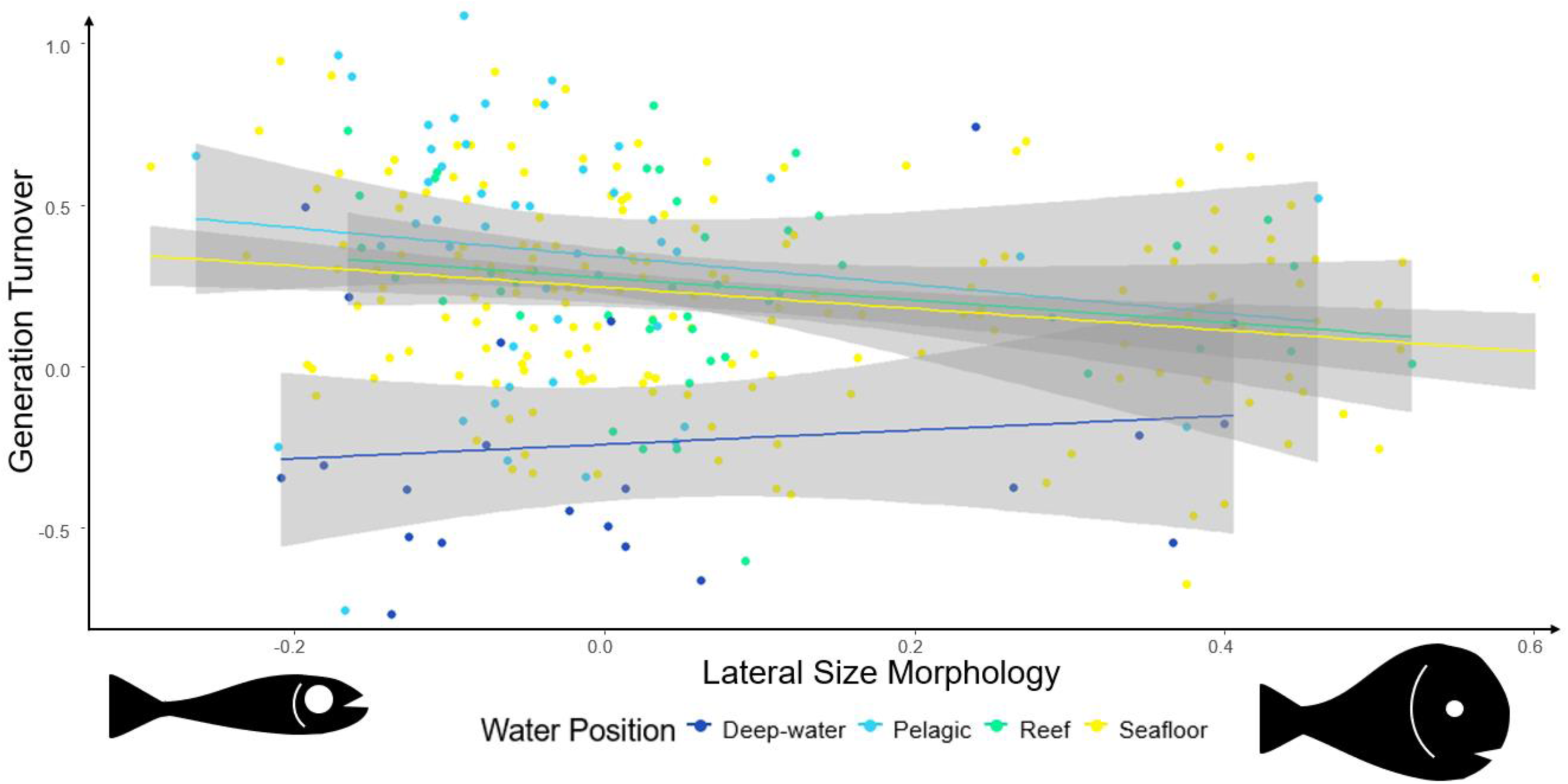
Lateral size morphology predicts generation turnover across different water positions. Scatter plot showing how lateral size the principal component (PC) axis lateral size morphology predicts the PC axis generation turnover differently depending on species position in the water column (grey areas denote 95% confidence intervals). The fish shadows indicate the scale for lateral size morphology from an elongated form with larger relative eye size and high oral gape position (left) to a rounder form with smaller eyes relative to the head and a low oral gape position (right).

**Figure 3:**
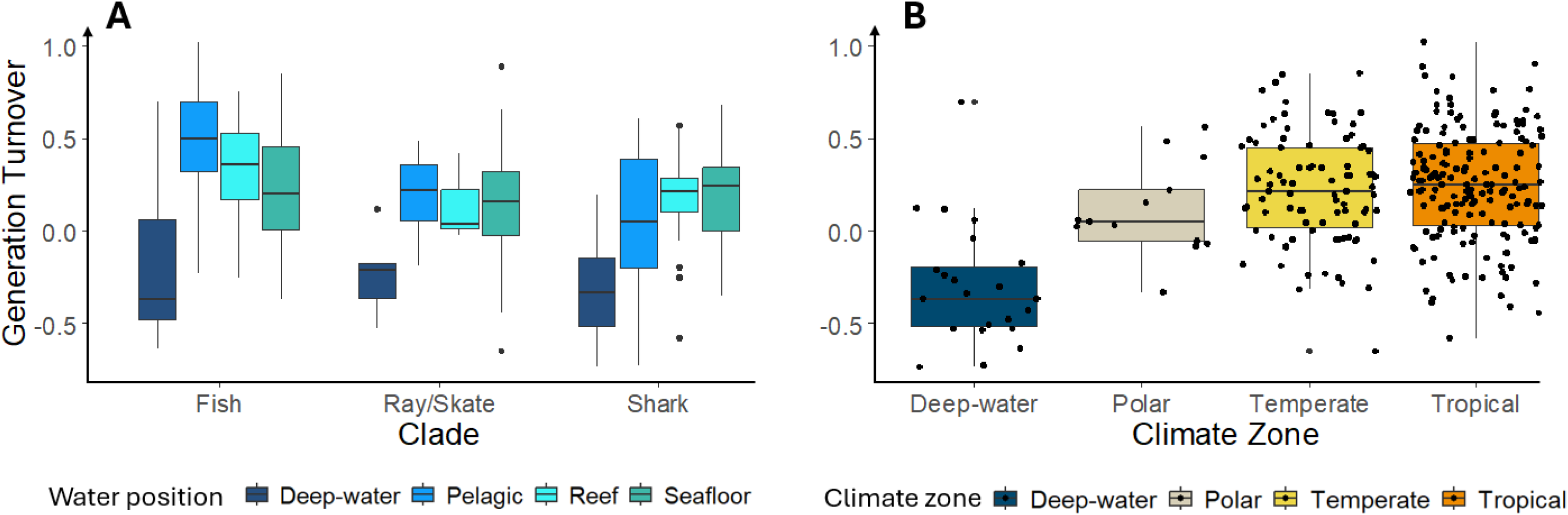
Generation turnover predicted by water position interacting with clade and climate zone. **A**: Boxplots showing how generation turnover principal component (PC) scores vary between ray-finned fish, rays/skates and sharks across different water positions (black dots are outlier species). **B**: Boxplots showing how generation turnover PC scores changes between deep-water, polar, temperate and tropical climate zones (black dots are species). In both A and B, vertical lines represent the range and horizontal lines of the boxplots are the medians.

### 3.3 Does morphology predict reproductive output?

If offspring production depends on energetic acquisition and allocation strategies that are shaped by foraging mode, locomotor performance and predator–prey interactions, we tested whether composite morphological axes linked to these performance traits predicted reproductive output across fish clades. In partial support of this prediction, the interaction between clade and lateral size morphology was statistically significantly associated with reproductive output LH-pPC scores (F_2, 274_=3.1796, p=0.0431). As lateral size morphology increased, reproductive output decreased in the ray-finned fish whereas it increased for the elasmobranchs (Figure 4A). The interaction between clade and locomotory morphology also statistically significantly predicted reproductive output (F_2, 274_=5.5527, p=0.0043). As locomotory morphology increased, reproductive output decreased in the ray-finned fish and increased in the elasmobranchs (Figure 4B). Positioning morphology statistically significantly predicted reproductive output (F_1, 274_=6.6295, p=0.01056). As positioning morphology increased, vertical eye and pectoral fin positions were lower and reproductive output decreased across all species (Figure 4C). Clade was also an independent predictor (F_2, 274_=15.586, p<0.001): ray-finned fish had the highest average reproductive output, followed by rays/skates and then sharks. No other interactions predicted reproductive output: climate zone × positioning morphology (F_2, 243_=0.7296, p=0.4832), climate zone × locomotory morphology (F_3, 245_=2.5782, p=0.0543), climate zone × lateral size morphology (F_3, 248_=0.8445, p=0.4707), water position × positioning morphology (F_3, 251_=2.3392, p=0.0739), water position × locomotory morphology (F_3, 254_=2.4261, p=0.06608), water position × lateral size morphology (F_3, 257_=0.5888, p=0.6229), clade × positioning morphology (F_2, 260_=0.5617, p=0.571), clade × water position (F_4, 262_=1.4914, p=0.1813), and clade × climate zone (F_6, 268_=0.3983, p=0.8799). The following single factors were retained but were not statistically significant predictors of reproductive output: positioning morphology (F_1, 274_=6.6295, p=0.0156), climate zone (F_3, 274_=1.8734, p=0.1343), clade (F_2, 274_=15.586, p<0.001), and water position (F_3, 274_=0.3042, p=0.8224). Lateral size morphology and locomotory morphology could not be reported on with ANOVA tests because the model became oversaturated. Instead, we report their test statistics from the final model: lateral size morphology (estimate=-0.3667, std. error=0.2798, t=-1.311, p=0.1910), locomotory morphology (estimate=-0.4356, std. error=0.2326, t=-1.873, p=0.0622).

**Figure 4:**
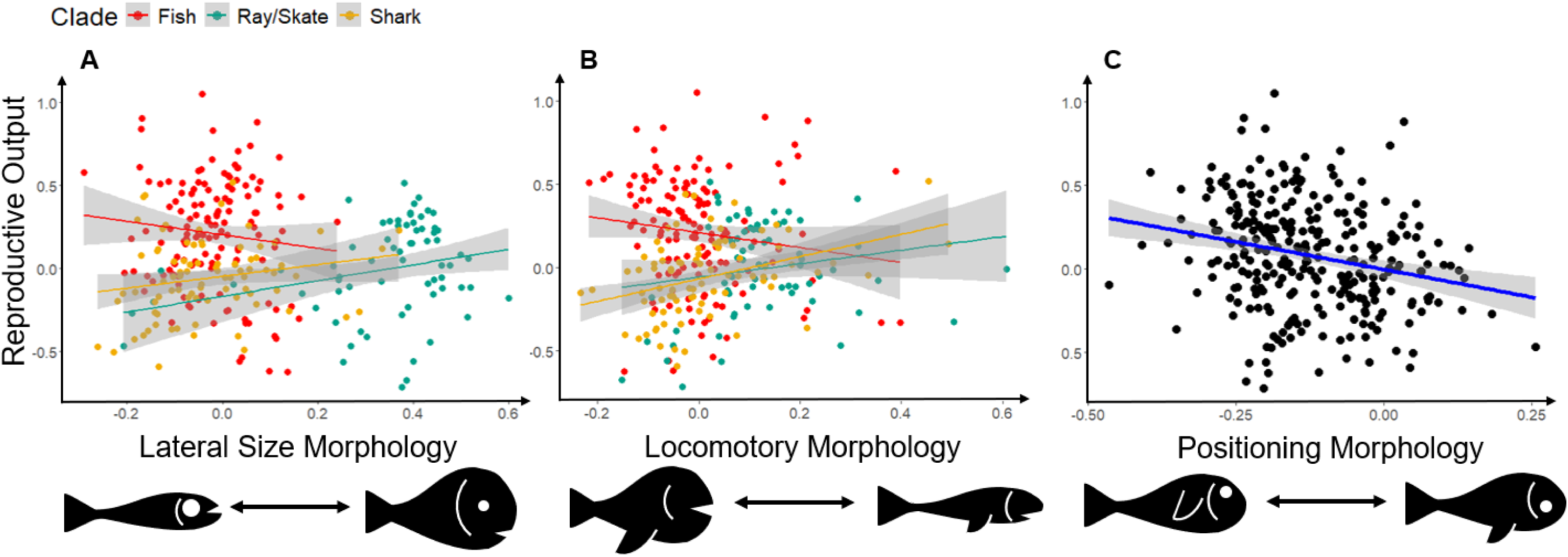
Composite morphological traits predict reproductive output. Scatter graphs showing how each of the morphological principal component (PC) axes scores predict reproductive output either interacting with clade (**A**, **B**) or independently (**C**). Each morphological axis is exemplified with fish shadows showing differences in body elongation, relative eye size, oral gape position, pectoral fin size and pectoral fin position. The linear relationships are shown by coloured lines which match the different clades (red=ray-finned fish, green=rays/skates, yellow=sharks), or are blue if no interactions were significant, and the grey shaded areas represent 95% confidence intervals.

## 4 DISCUSSION

Here, we tested the hypothesis that fish morphological traits linked to swimming behaviour, food acquisition and predator avoidance predict major life-history axes via shared performance trade-offs. Especially the consistent association between lateral size morphology and generation turnover across environments supports our hypothesis and illustrates how hydrodynamic trade-offs translate into demographic strategies.

### 4.1 Lateral size morphology as a predictor of generation turnover and fish performance across water positions

There is weak evidence to suggest that lateral size morphology is linked to generation turnover, dependent on a species’ position in the water column. In stable environments, generation time is often used as a measure of a species’ position along the slow–fast life-history continuum (Charlesworth, 1994; Heppell, Caswell & Crowder, 2000; Oli & Dobson, 2003; Gaillard et al., 2005; Stahl & Oli, 2006). Here, however, we found that a deeper body, smaller eyes relative to head depth, and a lower mouth position has an association with higher generation turnover in deep-water habitats, but lower generation turnover in pelagic, reef, and seafloor habitats. These results clearly support the ever-growing evidence that life history speed is mediated by ecological contexts, water depth in this case (Figure 3). The weak association observed here should therefore not be interpreted as evidence against a morphology–life history link. Rather, when considered alongside empirical studies documenting contrasting body morphologies across water depths, our results provide preliminary support for a common ecological driver linking morphology and life-history strategy. Testing this hypothesis within more closely related taxa or using more targeted functional traits may reveal stronger relationships than are apparent at this broad comparative scale (Kelly et al. 2021; Webb, 1984). Selection on ecological function can create consistent evolutionary associations between form and life-history strategy, because performance mediates survival, growth and reproduction (Clements and Ozgul 2016; Laughlin et al. 2020).

But what ecological function could drive the association between morphology and generation turnover? In pelagic, reef, and seafloor habitats, lower generation turnover may have an association with deeper bodies, smaller eyes and lower mouths. Although laterally compressed bodies incur greater drag, they enhance manoeuvrability, conferring advantages in predator evasion and prey capture (Howe, 2022), and can deter gape-limited predators (Brönmark & Miner, 1992; Nilsson, Brönmark & Pettersson, 1995). These benefits may lead to longer lifespans and thus slower generation turnover. In contrast, deep-water species show the opposite trend: lower generation turnover is linked to more elongate bodies, larger eyes and higher mouths. Below 200m, low light and high pressure select for energy-efficient, elongate morphologies adapted for slow, steady swimming (Seibel, Thuesen & Childress, 2000; Tytell et al., 2010; Neat & Campbell, 2013). Larger eyes enhance light sensitivity, improving detection of prey, predators, or conspecifics (Land, 1981; Myers et al., 2019), without the visibility costs that such traits can impose in illuminated habitats (Andersson, Scharnweber & Eklöv, 2024). Thus, opposing relationships between morphology and generation turnover arise because different depths impose contrasting selective pressures—balancing hydrodynamics, energy efficiency, and predator–prey interactions. Drawing these together may require testing different combinations and numbers of traits, focusing on those most closely influenced by the highlighted pressures, to reduce the noise found in the results here which may be obscuring the relationship between morphology and life history.

### 4.2 Morphology as a predictor of reproductive output and fish performance across clades

Similar to the links between morphology and generation time, the question is what ecological functions shape how morphology connects to reproductive output across different clades? Along the lateral size morphology axis, sharks and ray-finned fish with deeper bodies, lower mouths and smaller eyes occupy the higher end of the spectrum. This morphology is characteristic of demersal feeders, where enhanced manoeuvrability is advantageous in clustered habitats and large eyes are less critical because prey are often sedentary benthic invertebrates (Linsey & Collin, 2007; Gleiss, Potvin & Goldbogen, 2017; Howe, 2022). At the opposite end of the axis are sharks and ray-finned fish with elongated bodies and larger eyes positioned in shallower heads: a morphology typical of pelagic hunters, which benefit from reduced drag and improved visual tracking of mobile prey such as fish and cephalopods (Linsey & Collin, 2007; Gleiss, Potvin & Goldbogen, 2017). The positioning of rays and skates along this same axis reflects intra-clade differences: skates, with their shorter tails and reduced body elongation, plot higher than rays. Benthic-associated sharks and skates tend to exhibit higher reproductive output than pelagic-associated species because they are primarily oviparous (egg-laying), whereas many pelagic elasmobranchs are viviparous (live-bearing) (Katona et al., 2023). Consistent with the well-known trade-off between litter size and offspring size (Hussey, 2009; Sibly et al., 2018; Katona et al., 2023), oviparous elasmobranchs produce more but smaller offspring, while viviparous species produce fewer but larger ones. In pelagic systems, larger offspring have a greater chance of survival because of intense predation and limited refuges (Hussey, 2009; Sibly et al., 2018). This trade-off between survival/growth and reproduction explains the lower reproductive output of pelagic elasmobranchs. However, ray-finned fish show the opposite pattern: pelagic-associated species display higher reproductive outputs than benthic-associated ones. These species maximise recruitment success via broadcast spawning, releasing vast numbers of small eggs to offset high predation and resource limitation in the pelagic zone (Elgar, 1990; Winemiller & Rose, 1993; Sibly et al., 2018). Thus, the lateral morphology axis appears to capture a shared performance trade-off between offspring size and litter size that transcends clades, even though those trade-offs are realised through different reproductive tactics.

The link between morphology and reproductive output observed here is indirect, mediated through established trade-offs among growth, survival and reproduction. While the lateral size axis clearly reflects this pattern, relationships along the locomotory and positioning axes were weaker or reversed. For example, elongation along the locomotory axis did not consistently predict lower reproductive output, suggesting that different trade-offs between swimming behaviour and fecundity operate across species. Likewise, the relationship between vertical eye and pectoral-fin position and reproductive output hints at behavioural or ecological trade-offs related to prey and predator positioning in the water column, but these mechanisms remain speculative. One likely reason for this is that the morphological traits used in this study are functionally related to swimming, foraging and predator–prey interactions, but only indirectly to reproduction. Incorporating explicitly reproductive morphological traits (e.g. gonad size, oviduct morphology, clasper dimensions) could clarify these relationships, though such traits are often internal or not visible from lateral images. A more practical avenue may be to include additional life-history traits that capture reproductive trade-offs directly, such as offspring size, egg size or degree of parental care (Hussey, 2009; Sibly et al., 2018). Integrating these would better define the morphological correlates of reproductive strategy and improve the predictive power of morphology for life-history and population performance models.

## 5 CONCLUSION

Ultimately, this study provides proof of concept that morphology—particularly lateral size morphology tied to hydrodynamic constraints—can predict key life history axes and elements of population performance across diverse fish taxa. Body elongation, relative eye size and oral gape position emerged as strong correlates of generation turnover, linking water column position and swimming behaviour to life history strategy. These findings suggest that hydrodynamic forces may act as a unifying selective pressure shaping both morphology and demography in aquatic systems. As fish populations face accelerating pressures from overfishing, climate change and habitat loss, developing robust morphological proxies for life history will be essential. Our analyses represent a proof of concept rather than a predictive demographic framework. The life-history traits were derived under standardised, near-optimal conditions to reveal broad comparative patterns while minimising environmental variation among species. Although estimates based on maximum longevity should not be interpreted as realised demographic rates, they provide a consistent basis for identifying candidate morphology–life history relationships that can now be tested using species-specific demographic data and natural populations. Identifying a suite of externally measurable traits that reliably predict population-level traits would allow life history frameworks to be applied to many more data-deficient species—providing a scalable pathway to anticipate and mitigate biodiversity loss in changing aquatic ecosystems.

## ETHICAL APPROVAL

The work used published data on vertebrates that do not need ethical permissions.

## STATEMENT ON INCLUSION

This study prioritises broad taxonomic and ecological representation by including 290 marine and freshwater fish species spanning multiple clades, water column positions and climate zones. While no human subjects were involved, we acknowledge the importance of diversity in scientific collaboration and data sources. Future work should continue to expand inclusion of understudied regions and species to ensure equitable representation in global ecological and conservation research.

## AUTHOR CONTRIBUTIONS

V.D. conceived the ideas and V.D. and I.M.S. designed the methodology; V.D. collected the morphology data, analysed the data and led the writing of the manuscript. Both authors contributed critically to the drafts and gave final approval for publication.

## ACKNOWLEDGEMENTS

The authors thank Sol Lucas for support with analysing the data. ChatGPT-5 was used for language editing, grammar refinement, reference formatting and the creation of the graphical abstract. All scientific interpretation, data analysis, conclusions and final content were reviewed and approved solely by the authors.

## DATA AVAILABILITY STATEMENT

Data and code used to analyse the data run the models in MATLAB and R, as well as the Appendices are available at Figshare: https://figshare.com/s/32350693789254b34ae4.

